# Multi-echo Quantitative Susceptibility Mapping: How to Combine Echoes for Accuracy and Precision at 3 T

**DOI:** 10.1101/2021.06.14.448385

**Authors:** Emma Biondetti, Anita Karsa, Francesco Grussu, Marco Battiston, Marios C. Yiannakas, David L. Thomas, Karin Shmueli

## Abstract

**Purpose:** To compare different multi-echo combination methods for MRI quantitative susceptibility mapping (QSM). Given the current lack of consensus, we aimed to elucidate how to optimally combine multi-echo gradient-recalled echo (GRE) signal phase information, either before or after applying Laplacian-base methods (LBMs) for phase unwrapping or background field removal.

**Methods:** Multi-echo GRE data were simulated in a numerical head phantom, and multiecho GRE images were acquired at 3 T in ten healthy volunteers. To enable image-based estimation of GRE signal noise, five volunteers were scanned twice in the same session without repositioning. Five QSM processing pipelines were designed: one applied nonlinear phase fitting over echo times (TEs) before LBMs; two applied LBMs to the TE-dependent phase and then combined multiple TEs via either TE-weighted or signal-to-noise ratio (SNR)-weighted averaging; two calculated TE-dependent susceptibility maps via either multi-step or single-step QSM and then combined multiple TEs via magnitude-weighted averaging. Results from different pipelines were compared using visual inspection; summary statistics of susceptibility in deep gray matter, white matter, and venous regions; phase noise maps (error propagation theory); and, in the healthy volunteers, regional fixed bias analysis (Bland-Altman) and regional differences between the means (nonparametric tests).

**Results:** Nonlinearly fitting the multi-echo phase over TEs before applying LBMs provided the highest regional accuracy of *χ* and the lowest phase noise propagation compared to averaging the LBM-processed TE-dependent phase. This result was especially pertinent in high-susceptibility venous regions.

**Conclusion:** For multi-echo QSM, we recommend combining the signal phase by nonlinear fitting before applying LBMs.

## Introduction

MRI quantitative susceptibility mapping (QSM) aims to determine the underlying spatial distribution of tissue magnetic susceptibility (*γ*) from gradient-recalled echo (GRE) phase data (*ϕ*):

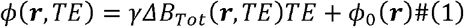

where ***r*** is a vector of image space coordinates, *γ* the proton gyromagnetic ratio, TE the echo time, Δ*B_Tot_* the *χ*-induced total field perturbation along the scanner’s z-axis, and *ϕ*_0_ the TE-independent phase offset at a nominal TE=0 ms.

For QSM, the acquired phase must be spatially (single-echo data) or spatiotemporally (multi-echo data) unwrapped to resolve 2*π* aliasing. The unwrapped *ϕ* is proportional to Δ*B_Tot_* (Equation 1), which is a combination of background (Δ*B_Bg_*) and local field contributions (Δ*B_Loc_*):

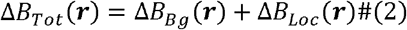

Δ*B_Bg_* are induced by the global geometry, air-tissue interfaces, and any field inhomogeneities. Δ*B_Loc_* reflect the tissue *χ* inside the region of interest (ROI), e.g., the brain. For QSM, Δ*B_Bg_* must be removed from Δ*B_Tot_*. The resulting Δ*B_Loc_* map is in the following relationship with *χ*:

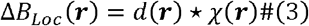

where *d* is the magnetic dipole and * denotes a spatially-dependent convolution. Based on Equation 3, the local distribution of tissue *χ*, i.e., the QSM map, is calculated by solving an ill-posed Δ*B_Loc_*-to-*χ* problem.

Recently, the QSM community critically reviewed how to best perform phase unwrapping (1) and Δ*B_Bg_* removal (2). Moreover, it promoted two challenges to compare algorithms for Δ*B_Loc_*-to-*χ* inversion, but a consensus has yet to be reached (3,4). A further open question toward QSM standardization is how and at which stage of the processing pipeline multi-echo data from different echoes should be combined. This question is relevant because phase unwrapping, Δ*B_Bg_* removal, or both, are often performed using Laplacian-based methods (LBMs) (1,2).

Laplacian phase unwrapping aims to calculate the 2π-aliasing-free phase as (5):

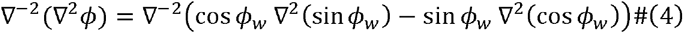

with ∇^2^ and ∇^−2^ the forward and inverse Laplace operators, and *ϕ_w_* the aliased phase.

Laplacian Δ*B_Bg_* removal relies on the harmonicity of Δ*B_Bg_* inside the ROI (i.e., ∇^2^Δ*B_Bg_* = 0, ***r*** ∈ *ROI*) and aims to solve this equation (2):

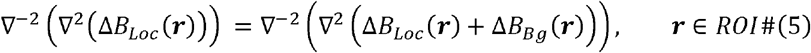

The inverse discrete Laplace operator is not well-defined and requires regularization, which is equivalent to spatially high pass filtering the phase or local field (2). However, the known varying frequency content at different TEs (6,7) could lead to different degrees of LBM-induced high-pass filtering at different echoes, alter the linearity of Equation 1, and thus introduce inaccuracies in the estimated Δ*B_Loc_* and *χ* maps. To investigate this issue, this study aimed to compare existing strategies for combining the multi-echo signal phase (see the Theory section for further details) when using LBMs for phase unwrapping or Δ*B_Bg_* removal.

Five processing pipelines for QSM were designed incorporating LBMs for both phase unwrapping and Δ*B_Bg_* removal and combining the signal from different TEs by fitting or averaging before or after applying LBMs. These pipelines were applied to both numerically simulated data and images acquired *in vivo*. Results from each pipeline were compared qualitatively by visual inspection and quantitatively via analysis of the regional *χ* bias and precision, and noise propagation.

## Theory

### Multi-echo combination

Previous studies employing multi-step reconstruction pipelines for QSM at 3 T have combined the signal from multiple echoes by either averaging (8–11) or fitting (12–15) before or after applying LBMs. For multi-echo combination in QSM, this study focuses on approaches based on weighted averaging (8–11) or complex nonlinear fitting (NLFit) (12,13) which outperform approaches based on unweighted averaging (16) or linear fitting (14).

#### Fitting

Multi-echo combination via nonlinear fitting (NLFit) has formulated the temporal evolution of the complex signal as a nonlinear least squares problem (12):

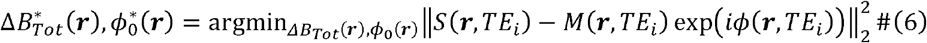

where *S* denotes the acquired complex signal, *M* the signal magnitude, *ϕ* the signal phase (Equation 1), and *TE_i_* the i-th echo time. This approach aims to mitigate noise in Δ*B_Tot_*, by correctly modeling as normally distributed the noise in the real and imaginary parts of the complex signal. Unlike weighted-averaging-based approaches, NLFit enables estimating *ϕ*_0_. Notably, the input phase to nonlinear fitting is minimally processed as it only requires to be temporally unwrapped, thus avoiding the application of LBMs before multi-echo combination.

#### Weighted averaging

Multi-echo combination (with *n* echoes) via weighted averaging has been performed using either TE-based weighting factors (8,9) or phase SNR-based weighting factors (10). TE-based weighted averaging (TE-wAvg) (8,9) accounts for the phase at shorter TEs being affected by larger noise levels than at longer TEs. TE-wAvg calculates a combined Δ*B_Tot_* as:

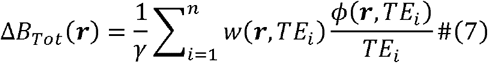

with weights equal to:

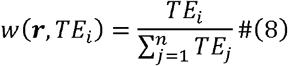

TE-wAvg requires temporal unwrapping of the input multi-echo phase, resulting in a combined Δ*B_Tot_*, which still contains Δ*B_Bg_* contributions.

SNR-based weighted averaging (SNR-wAvg) (10) accounts for different tissue types reaching optimal SNR at different TEs and calculates a combined local field map (Δ*B_Loc_*) as:

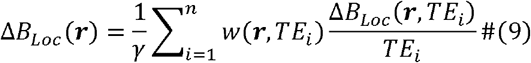

with weights equal to:

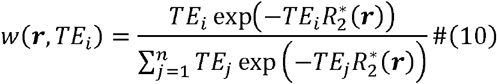

In Equation 10, 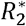 denotes the map of voxel-wise transverse relaxation rates, which is related to the signal magnitude *M* by:

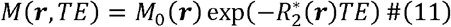

with *M*_0_ the initial transverse magnetization. SNR-wAvg requires performing temporal and spatial unwrapping as well as background field removal on the input multi-echo phase, resulting in a background-field-free field map.

Alternatively, a distinct *χ* map has been calculated at each TE and multi-echo combination of *χ* over time has been performed via weighted averaging using magnitude-based weighting factors (Susc-wAvg) (11). Based on the inversely proportionality of the phase noise and magnitude SNR (17), this method aims to improve the SNR of the combined *χ* map as:

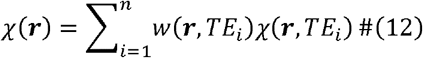

with weights equal to:

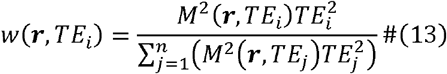

### Noise propagation

Previous studies using fitting (13) or SNR-wAvg (10) have calculated expressions for the noise in the total field map (σ(Δ*B_Tot_*)). For TE-wAvg or SNR-wAvg, expressions for σ(Δ*B_Tot_*) have not been calculated and were, therefore, derived here. For each multi-echo combination method, expressions for noise propagation from Δ*B_Tot_* to the corresponding Δ*B_Loc_* and *χ* images were also derived here.

Based on error propagation, the noise in Δ*B_Tot_* calculated using a linear least squares fitting approach (13) is (see Supporting Information):

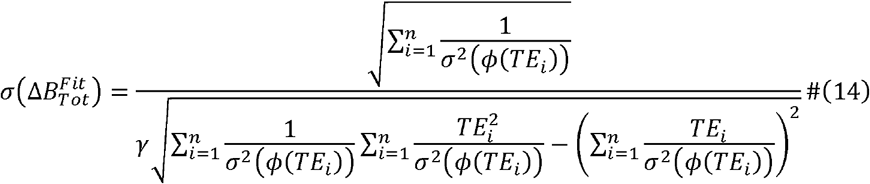

Equation 14 also corresponds to the a priori noise estimate found in nonlinear least squares fitting (12), thus 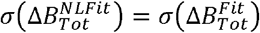.

Based on Equations 7–10, Δ*B_Tot_* calculated using TE-wAvg and Δ*B_Loc_* calculated using SNR-wAvg respectively have variances equal to:

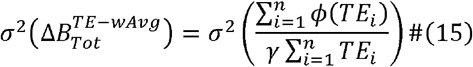

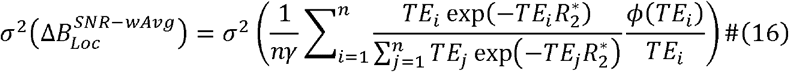

Assuming that noise in the single-echo phase is temporally uncorrelated and based on error propagation, the noise in Δ*B_Tot_*/Δ*B_Loc_* calculated using TE-wAvg/SNR-wAvg is respectively equal to:

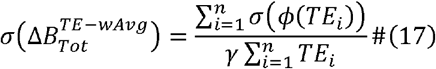

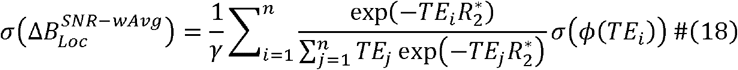

Equations 17 and 18 omit the ***r*** dependency, because they combine the multi-echo phase voxel by voxel.

#### Noise in the local field map

Based on error propagation and the orthogonality (18) of Δ*B_Loc_* and the Δ*B_Bg_* in the ROI, e.g., the brain:

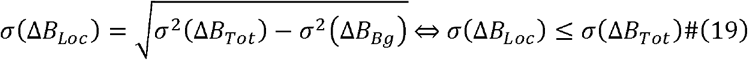

where ⇔ denotes an “if and only if” relationship, and σ(Δ*B_Loc_*) = σ(Δ*B_Tot_*) is the worst-case scenario.

#### Noise in the susceptibility map

Owing to the circular convolution theorem, the deconvolution operation in Equation 3 can be performed by point-wise division in the Fourier domain. Thus, if the regularized inverse dipole kernel in k-space 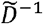 can be analytically derived independent of the Fourier transforms of *χ* or Δ*B_Loc_*, *σ*(*χ*) can be calculated as (see Supporting Information):

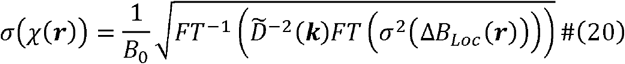

where *FT* and *FT*^−1^ respectively denote the direct and inverse Fourier transforms, and *k* denotes k-space coordinates. Deriving an analytical expression for 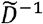 is possible, for example, when considering thresholded k-space division (TKD), or the Tikhonov-regularized minimal norm solution (13,19,20).

For weighted averaging of TE-dependent *χ* (Equations 12 and 13), based on phase error propagation over time and Equation 20, (*σΔB_Loc_*(*TE_i_*)) equals:

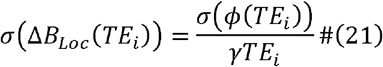

Based on Equations 12, 13, 20, and 21 (see Supporting Information), *σ*(*χ^susc-wAvgy^*) is the square root of:

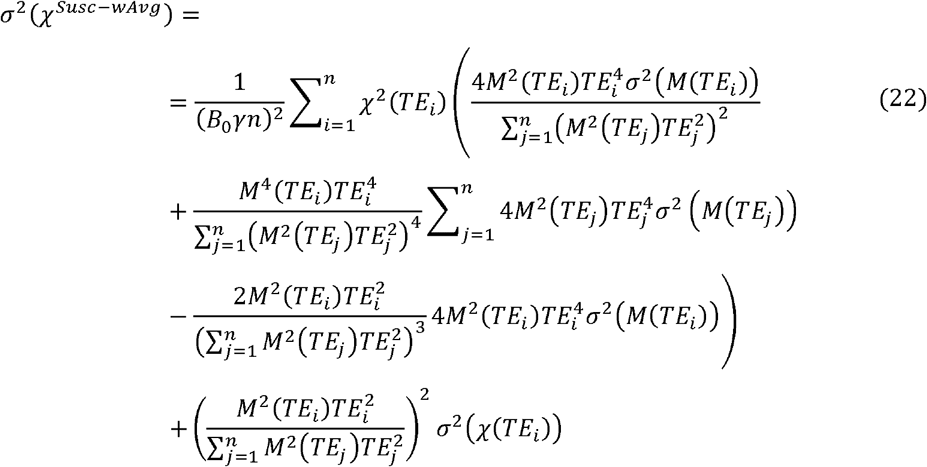

Notably, Equation 22 requires analytically describing *σ*(*χ*) (Equation 20).

## Methods

Where not otherwise stated, image analysis was performed on a 64-bit Windows 11 Pro operating system (Intel(R) Core(TM) i5-9400 CPU@2.90GHz processor; 16 GB RAM) using Matlab (R2021b, The MathWorks, Natick, MA). Preliminary versions of this study were presented at the 2016 and 2018 annual meetings of the International Society for Magnetic Resonance in Medicine (21,22).

### *In vivo* data acquisition

Multi-echo 3D GRE imaging of ten healthy volunteers (average age/age range: 26/22-30 years, five females) was performed in two centers (University College London Hospital and Queen Square Multiple Sclerosis Centre, University College London) equipped with the same 3 T MRI system (Philips Achieva, Philips Healthcare, Best, NL; 32-channel head coil). Five subjects were acquired in each center. All the volunteers provided written informed consent, and the local research ethics committees approved the experimental sessions. Images were acquired using a transverse orientation, field of view=240×240×144 mm^3^, voxel size=1-mm isotropic, flip angle=20□, repetition time=29 ms, five evenly spaced echoes (TE_1_TE spacing=3/5.4 ms), bandwidth=270 Hz/pixel, SENSE (23) factors=2/1.5, flyback gradients=on, no flow compensating gradients, total scan duration =04:37 min:s (24). Five subjects were scanned twice within the same session without repositioning to enable imagebased calculation of magnitude and phase SNRs.

### Data simulation from a numerical head phantom

To ensure the availability of ground-truth *χ* values against which to test the accuracy of QSM pipelines, a Zubal numerical head phantom was used (25) with the following regions of interest (ROIs): the caudate nucleus (CN), globus pallidus (GP), putamen (PU), thalamus (TH), superior sagittal sinus (SSS), grey and white matter (GM and WM), and cerebrospinal fluid (CSF). To match the acquisitions *in vivo*, the original 1.5-mm isotropic phantom was resampled to a 1-mm isotropic resolution with matrix size=384×384×192 voxels. Compared to our previous study (26), the numerical phantom was updated to achieve realistic regional means ± standard deviations (SDs) for *χ*, the proton density (*M*_0_), and the transverse relaxation rate 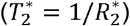 (see Supporting Information). Simulated multi-echo complex data were generated based on these ground-truth spatially variable *χ*, *M*_0_ and 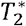 distributions, as:

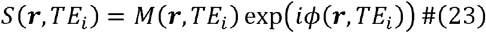

with *ϕ* and *M* respectively described by Equations 1 and 11, and TEs matched to the *in vivo* acquisitions. Random zero-mean Gaussian noise with a SD = 0.07 was added to the real and imaginary parts of the noise-free signal independently (17,26). The random noise matrix was regenerated at each TE.

### Data preprocessing

A brain mask was calculated for each subject by applying FSL BET (27,28) with robust brain center estimation (threshold=0.3) to the magnitude image at the longest TE. This choice of TE accounted for the greater amount of signal dropout near regions of high-*χ* gradients compared to shorter TEs.

A whole-brain mask for the Zubal phantom was calculated by applying FSL BET with robust brain center estimation (threshold=0.5) to the 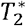 map of the numerical phantom (26).

### Processing pipelines for QSM

Five distinct processing pipelines (Figure 1) were applied to both the numerically simulated and the healthy volunteer data, and the time required to run each pipeline was measured using Matlab’s stopwatch timer. Three of these pipelines (NLFit, TE-wAvg, and SNR-wAvg) combined the phase across TEs at different stages before performing the Δ*B_Loc_*-to-*χ* inversion. Two other pipelines (Susc-wAvg and Susc-TGV-wAvg) first calculated a distinct *χ* map at each TE and then combined the *χ* maps. The following paragraphs describe each processing pipeline in detail.

**Figure 1.**
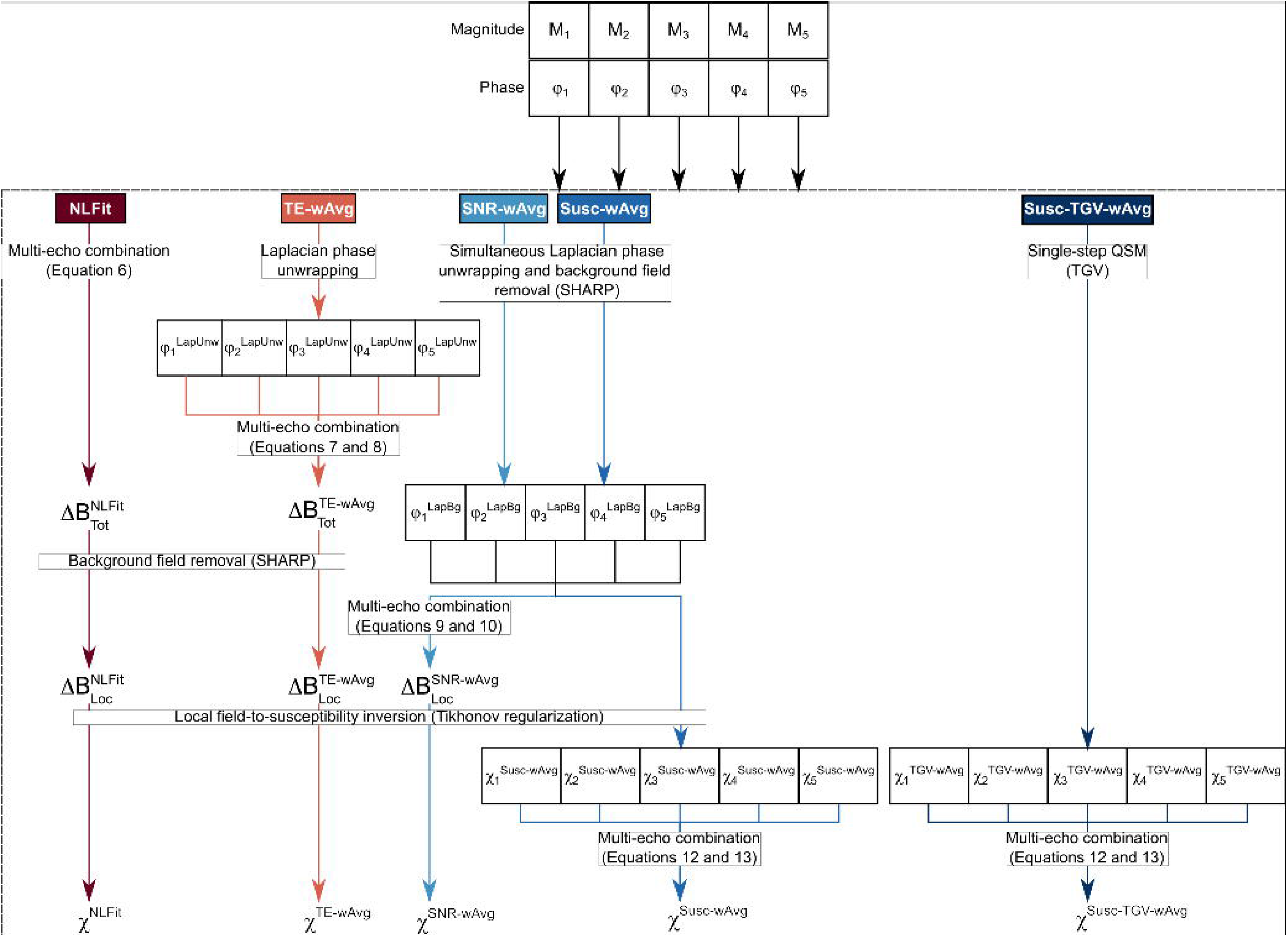
Processing pipelines for multi-echo QSM. For each multi-echo combination method (NLFit [nonlinear phase fitting], TE-wAvg [TE-weighted phase averaging], SNR-wAvg [SNR-weighted phase averaging], Susc-wAvg and Susc-TGV-wAvg [magnitude-weighted susceptibility averaging]) the processing steps are described as separate processing streams.

The NLFit pipeline (26) first combined the complex GRE signal by nonlinear fitting over TEs (12) using the Cornell QSM software package’s *Fit_ppm_complex* function (29). It then applied simultaneous spatial phase unwrapping and Δ*B_Bg_* removal using Sophisticated Harmonic Artifact Reduction for Phase data (SHARP) (30), a direct solver of Equation 5 (30). SHARP was chosen because it has been widely used in the literature on QSM and is both robust and numerically efficient (2). Moreover, in our recent study comparing multi-echo and TE-dependent QSM, a multi-echo pipeline incorporating SHARP gave highly accurate multiecho QSM values (26). SHARP was applied using the minimum-size 3-voxel isotropic 3D Laplacian kernel (30), a threshold for truncated singular value decomposition (tSVD) equal to 0.05, and a brain mask eroded by five voxels.

The TE-wAvg processing pipeline first applied Laplacian unwrapping to the phase at each TE using a threshold for tSVD equal to 10^-10^ (i.e., the default value in (29)). Second, it calculated Δ*B_Tot_* by averaging the unwrapped phase according to Equations 7 and 8 (8,9). Then, it calculated Δ*B_Loc_* by applying SHARP with the same parameters as in NLFit.

The SNR-wAvg processing pipeline first applied simultaneous phase unwrapping and Δ*B_Bg_* removal to the phase at each TE using SHARP with the same parameters as in NLFit. An 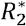 map was calculated by voxel-wise fitting Equation 11 using Matlab’s *nlinfit* function, where initial values for 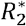 and *M*_0_ were calculated by linearly fitting the log-linearized version of Equation 11. The SNR-wAvg pipeline then calculated Δ*B_Loc_* by averaging the unwrapped and background-field-free phase according to Equations 9 and 10 (10).

In the NLFit, TE-wAvg, and SNR-wAvg pipelines, Δ*B_Loc_*-to-*χ* inversion was performed using Tikhonov regularization with correction for susceptibility underestimation and using the L-curve method to determine the optimal value for the regularization parameter (13,30,31). This inversion method was chosen because it is computationally efficient and substantially reduces streaking artifacts relative to TKD (32).

The Susc-wAvg processing pipeline calculated a distinct *χ* map at each TE by applying simultaneous phase unwrapping and Δ*B_Bg_* removal using SHARP as in NLFit and performing the Δ*B_Loc_*-to-*χ* inversion using Tikhonov regularization as in NLFit, TE-wAvg and SNR-wAvg. This pipeline then calculated a combined *χ* map according to Equations 12 and 13 (11).

The Susc-TGV-wAvg processing pipeline applied one-step TGV (33) to the phase at each TE, and then calculated a combined *χ* map as in Susc-wAvg. The TGV method was tested because it avoids stair-casing artifacts in the resulting *χ* map while correctly preserving structural borders (33). Moreover, in our recent study comparing multi-echo and TE-dependent QSM, TGV provided highly accurate TE-dependent QSM images (26). TGV (v1.0.0_20210629) was run in Neurodesk (v20220302, https://neurodesk.github.io/) with the default parameter values (*α*_1_, *α*_0_) = (0.0015,0.005), which are optimal for medical imaging applications (33).

### ROI segmentation in the healthy volunteer images

Regional *χ* values were compared within the simulated and *in vivo* data, similarly to our previous study (26). ROIs similar to those in the numerical phantom were segmented *in vivo*: the CN, GP, PU, TH, posterior corona radiata (PCR) as a WM ROI, and the straight sinus (StrS) as a venous ROI. Briefly, for each subject, the CN, GP, PU, TH and PCR were segmented based on the Eve *χ* atlas (34), whose GRE magnitude image was aligned to each subject’s fifth-echo magnitude image using NiftyReg (35,36) (TE_Eve_/TE_5_=24/24.6 ms). The quality of ROI alignment was assessed by visual inspection. The ITK-SNAP active contour segmentation tool (37) was used to segment the StrS over several slices based on the fifth-echo magnitude image, which presented the best contrast between the StrS and the surrounding brain tissue.

### Quantitative evaluation of the measured *χ*

In the numerical phantom simulations, each QSM pipeline’s performance relative to the ground truth was visually assessed by calculating a difference image between the corresponding Δ*B_Loc_/χ* map and 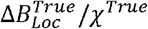. Here, 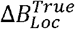 referred to the ground-truth local field calculated using the reference scan method (18,38), and *χ^True^* to the ground-truth magnetic susceptibility distribution with realistic regional means ± SDs of *χ* (Supporting Information Figure S1B). Means and SDs of *χ* were calculated for each pipeline in each ROI with *χ^True^* ≠ 0, i.e., the CN, GP, PU, TH, SSS, and WM. The root mean squared errors (RMSEs) of both Δ*B_Loc_* and *χ* relative to *χ^True^* were also calculated throughout the brain volume (26). Notably, RMSEs for Δ*B_Loc_* could not be calculated for the one-step Susc-TGV-wAvg pipeline.

In the volunteers, owing to the lack of a ground truth, representative susceptibility difference images were calculated relative to *χ^NLFit^*, because the NLFit pipeline performed multi-echo combination at the earliest possible stage, and relative to *χ^TE–wAvg^*, because the TE-wAvg pipeline had the lowest local field RMSE in the numerical phantom simulations (see the Results). Regional means and SDs of *χ* were calculated for each processing pipeline and compared against *χ* values in subjects of a similar age from the QSM literature. RMSEs could not be calculated because of the lack of a ground truth. For visualization purposes, the pooled averages and SDs were calculated (26) after verifying that all *intra*subject SDs of *χ* were larger than the *inter*subject SD of *χ*.

### Noise propagation maps

Only the healthy volunteers scanned twice were considered for this analysis. To enable image SNR calculation, in one healthy volunteer, five 20×20-voxel ROIs were drawn on a sagittal slice of the first-echo magnitude image (39), including both the GM and WM and excluding regions with artifacts induced by SENSE, motion, or flow. All five ROIs were applied across the other four volunteers by using rigid alignment transforms (NiftyReg (36)) between the first-echo magnitude images.

In each subject, ROI-based magnitude (*M_ROI_*), magnitude noise (σ*_ROI_*(*M*)) and phase noise values (σ*_ROI_*(*ϕ*)) were calculated based on the SNR difference method (40):

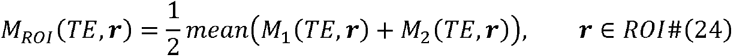

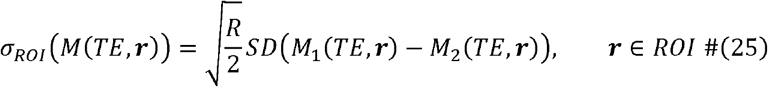

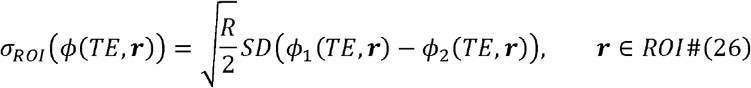

Here, *M*_1_/*ϕ*_1_ and *M*_2_/*ϕ*_2_ respectively denote the magnitude/phase images from the first and second scan, *R* = 3 is the 2D SENSE factor calculated by multiplying the SENSE factors applied along the two phase encoding directions (41). The values calculated based on Equations 24–26 were averaged across the five ROIs, to calculate summary values of magnitude, magnitude noise, and phase noise at each TE.

For both the numerical phantom simulations and the healthy volunteers, a phase noise σ(*ϕ*) map was calculated at each TE as (17,42):

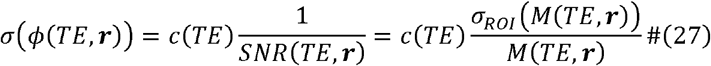

where *c*(*TE*) was a constant equal to 1 (by definition) in the numerical phantom simulations and equal to:

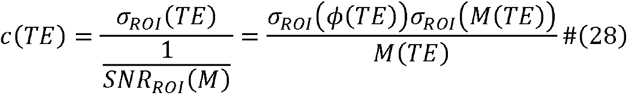

in the healthy volunteers. Notably, calculating phase noise analytically as in Equation 27 enabled the direct comparison of noise propagation between numerical simulations and data acquired *in vivo*.

For the NLFit, TE-wAvg and SNR-wAvg pipelines, the σ(Δ*B_Loc_*) map was calculated based on the multi-echo σ(*ϕ*) maps according to Equations 14 and 17–19. Then, the σ(*χ*) map was calculated according to Equation 20 with *B*_0_ = 3 T and the Tikhonov-regularized inverse magnetic dipole kernel with regularization parameter *α* (13):

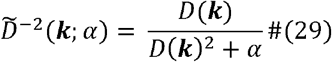

and correction for *χ* underestimation (30). For the Susc-wAvg pipeline, the σ(*χ*) map was calculated based on Equation 22. σ(*χ*) maps were not calculated for the Susc-TGV-wAvg pipeline, because TGV estimates *χ* iteratively (33) and an analytical expression for σ(*χ*) could not be derived.

To compare σ(*χ*) maps across pipelines, a line profile was traced in the same location of all σ(*χ*) images, and the σ(*χ*) value of each voxel along this profile was plotted. To compare the noise intensity and its variability between processing pipelines, the mean and SD of this representative line profile were also calculated.

### Statistical analysis

Statistical analyses were performed based on the healthy subject data. For each ROI and each pair of pipelines, Bland-Altman analysis of the average *χ* was used to assess if pairs of pipelines systematically produced different results. For each ROI, statistically significant differences between pipelines were tested by considering the corresponding distributions of average *χ* values across subjects. To assess whether to apply parametric paired *t*-tests or nonparametric sign tests, the normal distribution of the differences between paired *χ* values was assessed using the Shapiro-Wilk test. All statistical tests were two-tailed, and an uncorrected *p*-value < 0.05 was considered significant.

## Results

### Pooling of *χ* measurements

For each ROI and each processing pipeline, all *intra*subject SDs of *χ* were larger than the *inter*subject SD. Thus, pooled means and SDs were calculated (26).

### Performance of pipelines for multi-echo QSM

The Susc-wAvg and Susc-TGV-wAvg processing pipelines were the longest to run (Supporting Information Table S2), as they calculated a QSM map at each TE. The longer processing times required for the numerical phantom data were linked to the larger matrix size (384×384×192) compared to acquisitions *in vivo* (240×240×144).

In the numerical phantom, Figure 2 shows the ground-truth *χ* (A, G), the QSM images calculated by each processing pipeline (B-L), their difference relative to the ground truth (MV), and the RMSEs of *χ* throughout the brain volume (bottom row). Analogous results for Δ*B_Loc_* are shown in Supporting Information Figure S3. In the numerical phantom simulations, the *χ* map calculated using the NLFit pipeline had the largest RMSE (109.4%) followed, in decreasing order, by the SNR-wAvg (94.3%), TE-wAvg (93.5%), Susc-wAvg (91.5%) and Susc-TGV-wAvg pipelines (80.4%) (Figure 2, bottom row). The Δ*B_Loc_* map calculated using the Susc-wAvg pipeline had the largest RMSE (average across TEs: 85.0%) followed, in decreasing order, by the NLFit (81.2%), TE-wAvg and Susc-wAvg pipelines (both 71.8%).

**Figure 2.**
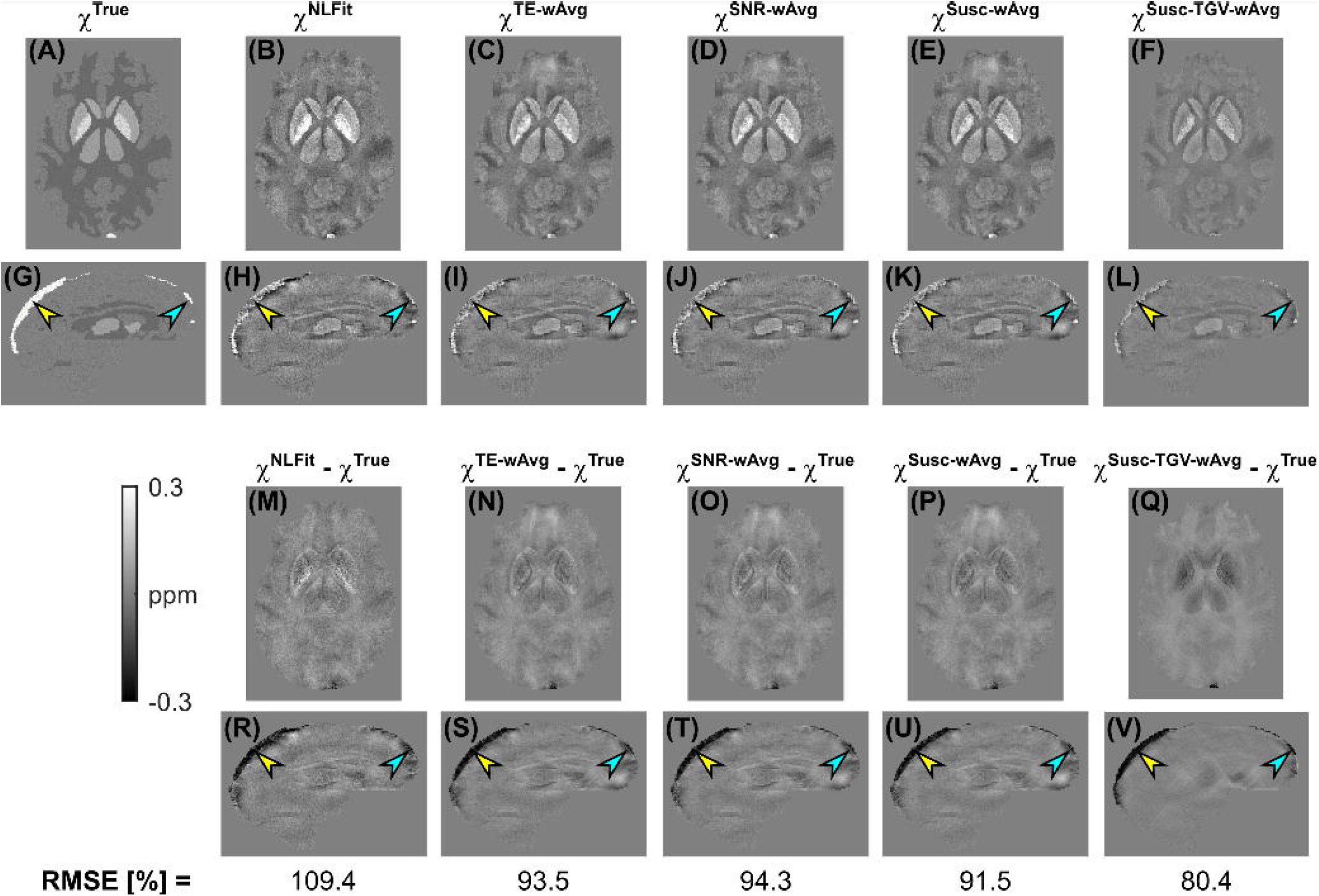
*χ* maps calculated using distinct multi-echo combination methods in the numerical phantom simulations. The same transverse and sagittal slices are shown for the ground-truth susceptibility map **(A, G)**, and for the susceptibility maps calculated using NLFit **(B, H)**, TE-wAvg **(C, I)**, SNR-wAvg **(D, J)**, Susc-wAvg **(E, K)**, and Susc-TGV-wAvg (**F, L)**. The figure also shows the difference between each susceptibility map and the ground truth **(M-V)**. The bottom row shows the root mean squared errors (RMSEs) of *χ* for each pipeline. In all the sagittal images **(G-L, R-V)**, the yellow and blue arrowhead respectively point at the same posterior and anterior locations in the superior sagittal sinus.

In the numerical phantom and for each processing pipeline, Figure 3A shows the regional means and SDs of *χ*. The error between the calculated and ground-truth *χ* appeared similar for all processing pipelines, although slightly larger SDs were always observed for the NLFit pipeline (Figures 2M-V and 3A). The SSS, which was the structure with the largest *χ^True^*, showed the largest susceptibility errors for all processing pipelines (arrowheads in Figures 2G-L, R-V, and Figure 3A).

**Figure 3.**
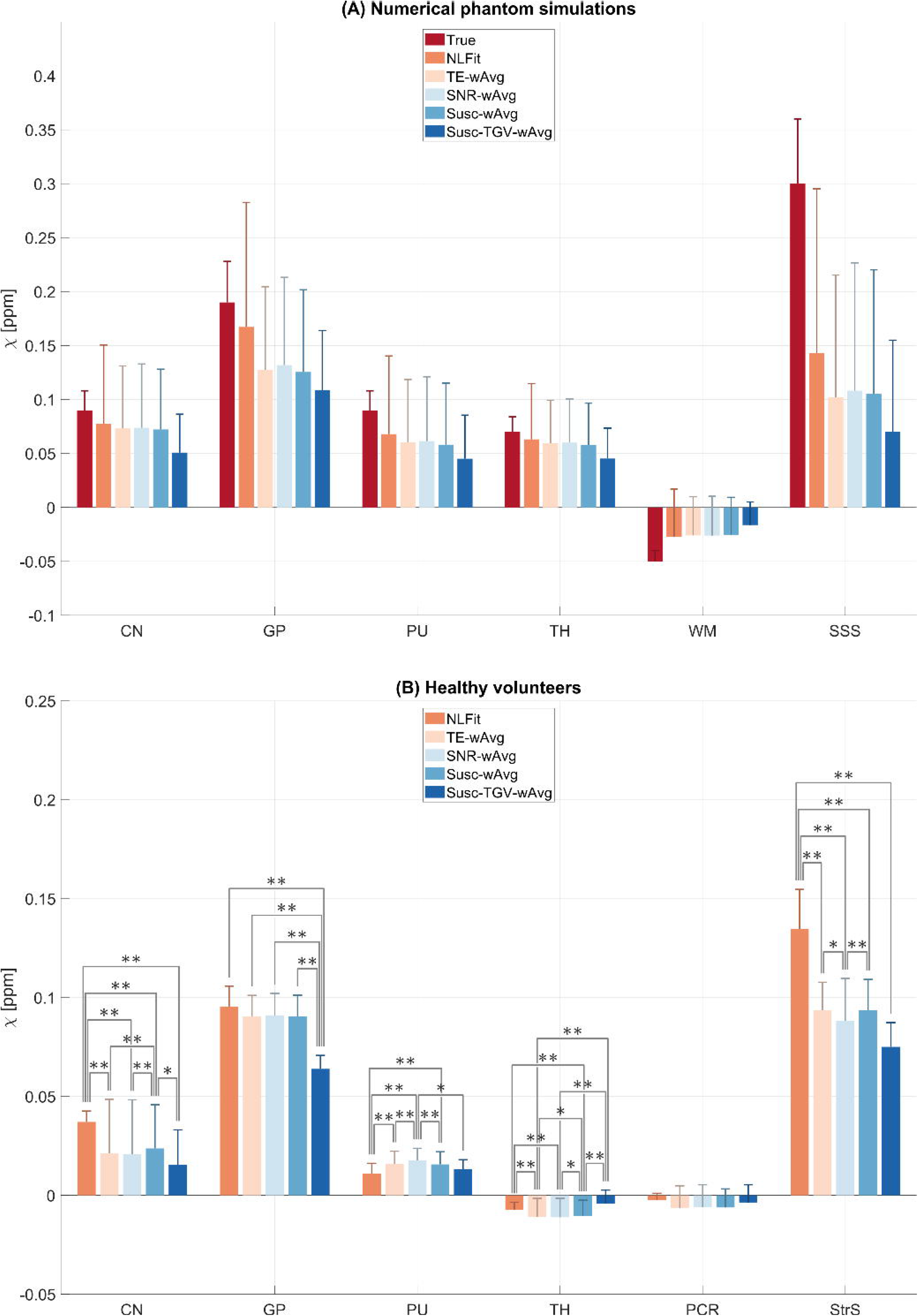
Means and SDs of *χ* in the phantom and healthy volunteer ROIs. The means and SDs (error bars) of *χ* are shown in each ROI of the numerically simulated **(A)** and pooled healthy volunteer data **(B)** for each processing pipeline. In the numerical phantom the ground-truth *χ* is also shown. In the healthy volunteers, significant differences between pairs of pipelines are denoted using the symbols * (*p*-value < 0.05) and ** (*p*-value < 0.01). Abbreviations: CN, caudate nucleus; GP, globus pallidus; PCR, posterior corona radiata; PU, putamen; StrS, straight sinus; TH, thalamus; WM, white matter; SSS, superior sagittal sinus.

For one representative volunteer, Figures 4A-J show the susceptibility images calculated using each processing pipeline. Additionally, differences images are shown relative to the *χ^NLFit^* (K-R) and the *χ^TE–wAvg^* maps (S-Z). Here, susceptibility differences between processing pipelines were most prominent in the StrS (arrowheads in Figures 4F-J, O-R, W-Z). In the healthy volunteers, Figure 3B shows the pooled regional means and SDs of *χ* calculated by each processing pipeline. The average *χ* measured in the deep-GM ROIs and in the PCR had values within the ranges reported by previous studies (34,43–47): 0.01–0.13 ppm for the CN, 0.06-0.29 ppm for the GP, 0.02–0.14 ppm for the PU, −0.02–0.08 ppm for the TH, and −0.06–0.03 ppm for the PCR. In the StrS, only *χ^NLEit^* had an average value close to the previously reported range for venous blood, namely, 0.17–0.58 ppm (32,46,48,49) (Figure 3B).

**Figure 4.**
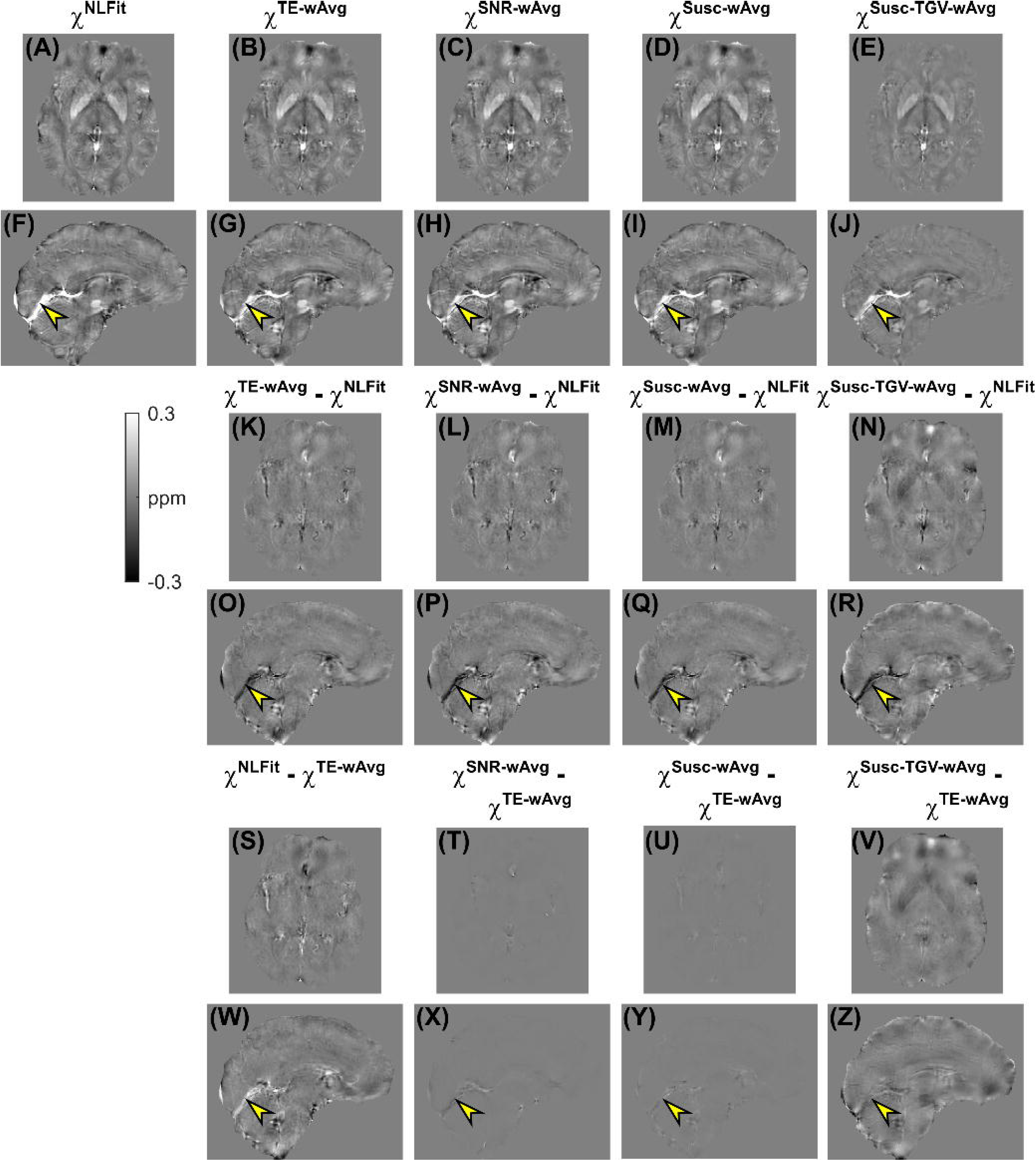
*χ* maps calculated using distinct multi-echo combination methods in a representative healthy volunteer. The same transverse and sagittal slices are shown for the susceptibility maps calculated using NLFit **(A, F)** [nonlinear phase fitting], TE-wAvg **(B, G)** [TE-weighted phase averaging], SNR-wAvg **(C, H)** [SNR-weighted phase averaging], Susc-wAvg **(D, I)** and Susc-TGV-wAvg **(E, J)** [magnitude-weighted susceptibility averaging]. The figure also shows the differences between the TE-wAvg, SNR-wAvg, Susc-wAvg and Susc-TGV-wAvg maps and the NLFit map **(K-R)**, and the differences between the NLFit, SNR-wAvg, Susc-wAvg and Susc-TGV-wAvg maps and the TE-wAvg map **(S-Z)**. In all the sagittal images **(F-J, O-R, W-Z)**, the yellow arrowheads point at the same location in the StrS.

In the CN, GP, and venous ROIs, the regional *χ* measured *in vivo* had a relative accuracy across pipelines similar to the numerical phantom simulations: the NLFit and Susc-TGV-wAvg pipelines respectively provided the highest and lowest means of *χ*, whereas the TE-wAvg, SNR-wAvg, and Susc-wAvg pipelines provided intermediate values (Figure 3). Slightly different trends were observed in the PU, TH, and WM ROIs, where, *in vivo*, the NLFit and Susc-TGV-wAvg pipelines provided the lowest (absolute) means of *χ*, and the TE-wAvg, SNR-wAvg, and Susc-wAvg pipelines provided higher values (Figure 3B). In the numerical phantom simulations, the NLFit pipeline always had the largest SD of *χ* (Figure 3A). In contrast, *in vivo*, the NLFit pipeline had the smallest SD of *χ* in the CN, PU, TH, and PCR, and SDs of *χ* comparable to other pipelines in the GP and StrS (Figure 3B).

### Phase noise propagation into the *χ* maps

Figures 5 and 6 show the estimated σ(*χ*) maps and profiles in the numerical phantom simulations and a representative healthy subject, respectively. All σ(*χ*) images contained some degree of streaking artifacts, especially *in vivo*, as these are a manifestation of error propagation from Δ*B_Loc_* to *χ* caused by the dipole kernel null space (20). In the phantom, the NLFit and Susc-wAvg pipelines had the σ(*χ*) line profiles with the lowest means and SDs (Figures 5A, D, E, H, I). The NLFit and SNR-wAvg pipelines always resulted in the lowest streaking artifacts burden (Figures 5A, E, C, G, I and 6A, E, C, G, I). Streaking artifacts were more severe in the TE-wAvg and Susc-wAvg pipelines especially near high-*χ* venous structures.

**Figure 5.**
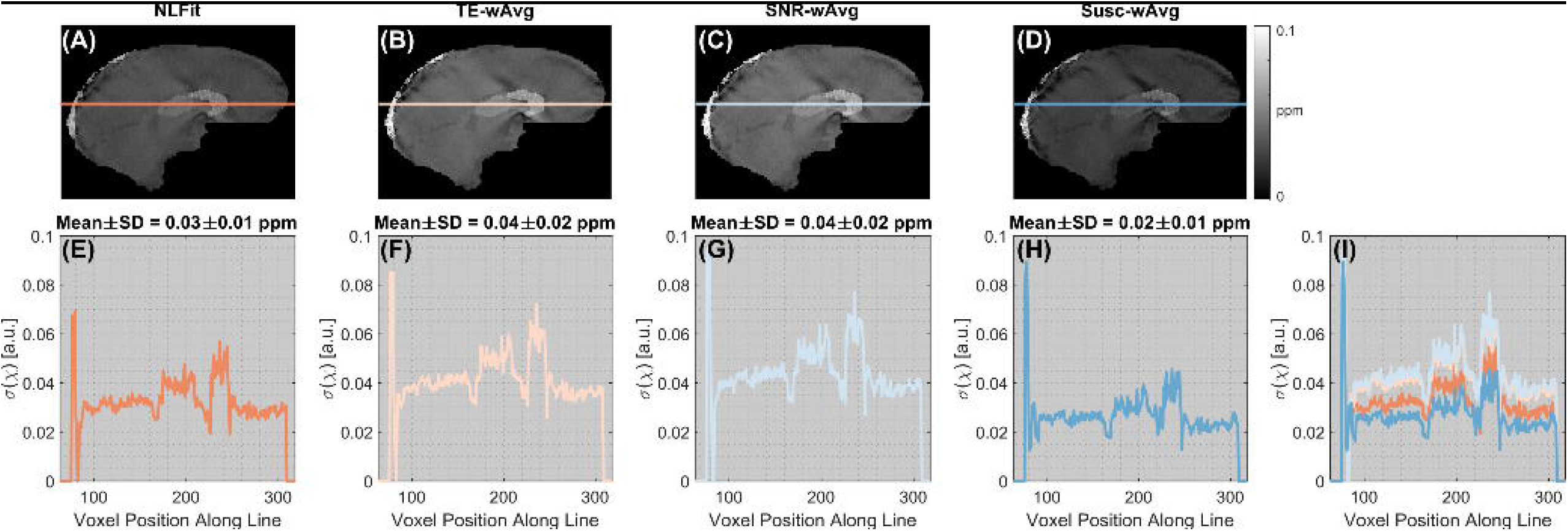
Susceptibility noise profiles in the numerical phantom simulations. The same sagittal slice is shown for susceptibility noise (σ(*χ*)) images calculated using the NLFit, TE-wAvg, SNR-wAvg and Susc-wAvg pipelines **(A-D)**. The susceptibility noise is plotted for a line profile traced on the σ(*χ*) images **(E-H)** including the high-*χ* SSS. All line profiles are also shown combined in the same plot **(I)**. The mean and the standard deviation of σ(*χ*) are shown for each line profile.

**Figure 6.**
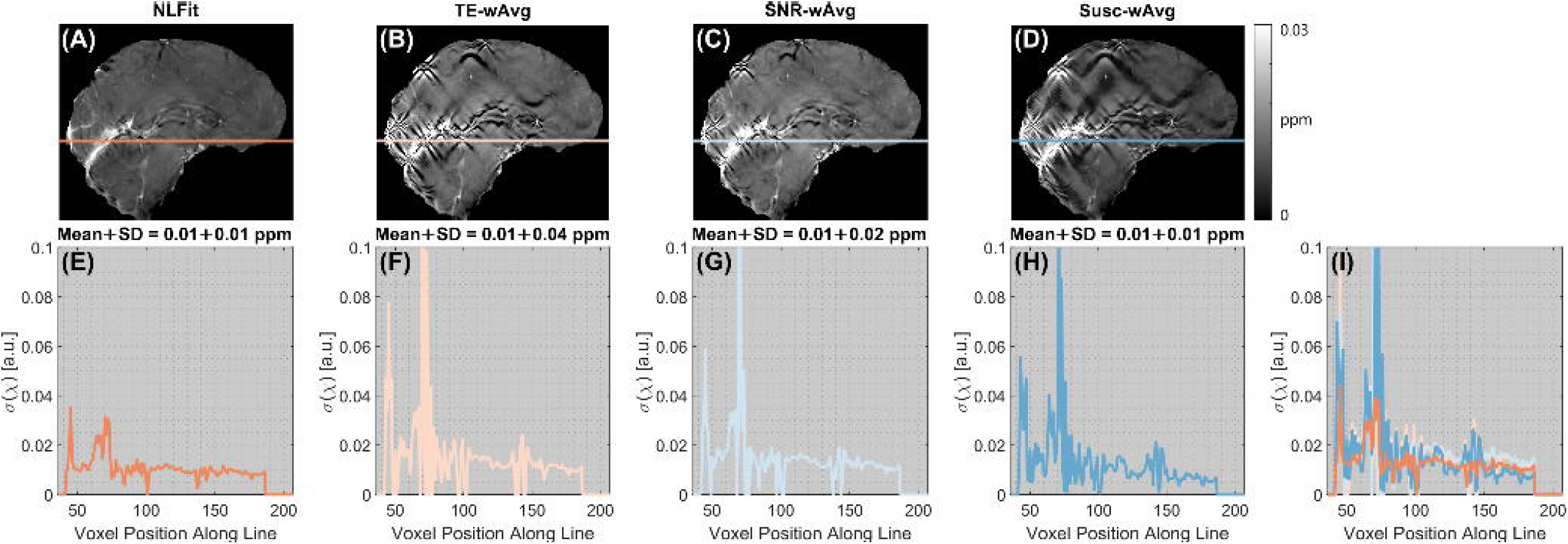
Susceptibility noise profiles in a representative healthy volunteer. The same sagittal slice is shown for susceptibility noise (σ(*χ*)) images calculated using the NLFit, TE-wAvg, SNR-wAvg and Susc-wAvg pipelines **(A-D)**. The susceptibility noise is plotted for a line profile traced on the σ(*χ*) images **(E-H)** including the high-*χ* StrS. All line profiles are also shown combined in the same plot **(I)**. The mean and the standard deviation of σ(*χ*) are shown for each line profile.

### Statistical Analysis

In the healthy volunteers, for all processing pipelines and ROIs, the Shapiro-Wilk test always rejected the hypothesis of normally distributed paired differences of *χ*. Therefore, pairwise comparisons between pipelines were always evaluated using the nonparametric sign test. Significant differences between pipelines are shown in Figure 3B, whereas the between-pipeline biases are shown in Figure 7. A lower threshold equal to |0.01| ppm (|·| denotes the absolute value) was chosen because, below this level, *inter*pipeline differences cannot be disentangled from the *intra*pipeline variability (i.e., the regional SD).

**Figure 7.**
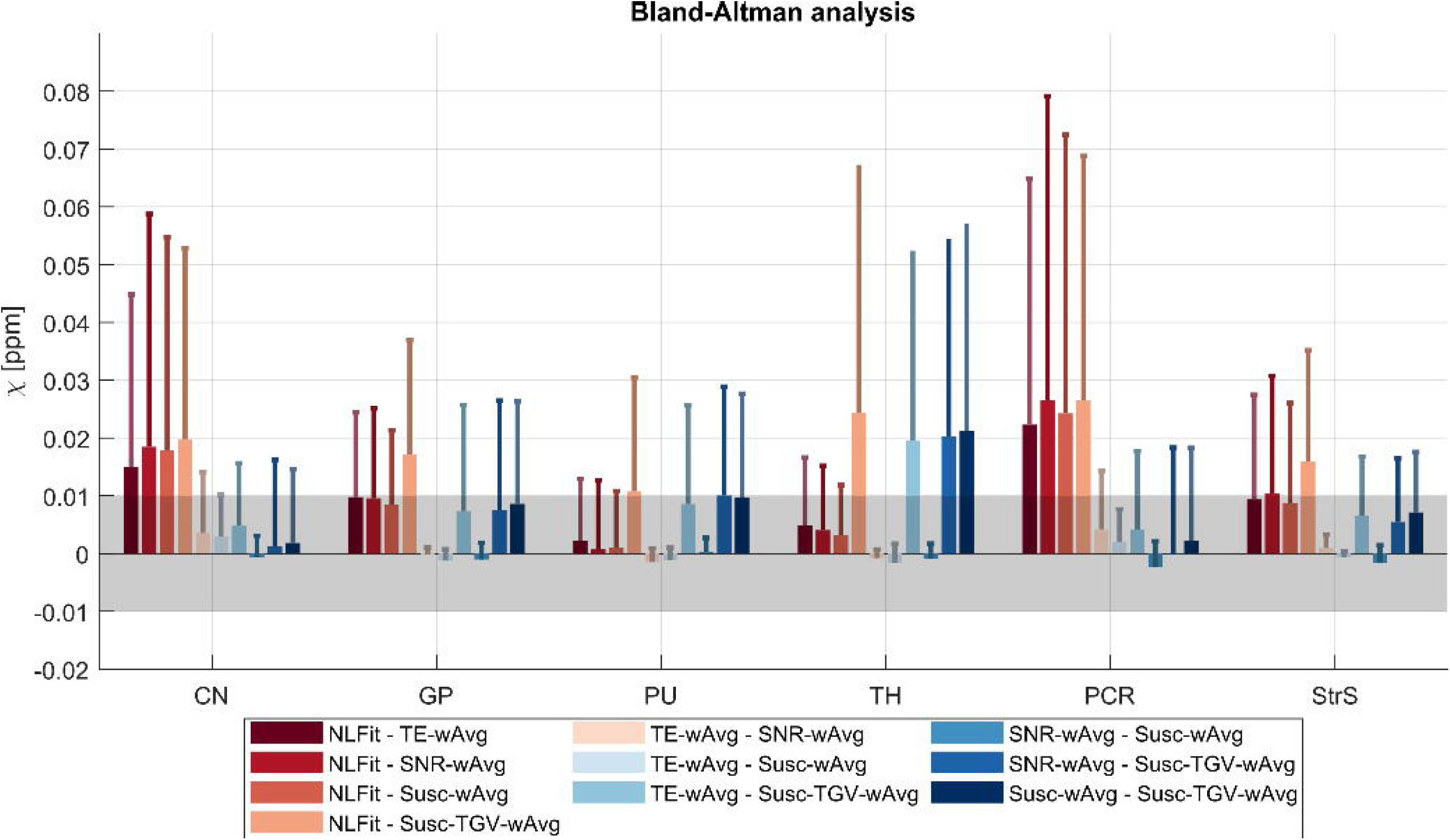
Bias between multi-echo pipelines for QSM in each ROI. The mean and SDs (error bars) of the bias are shown in each healthy volunteer ROI for all pairs of multi-echo processing pipelines. The grey band denotes the [−0.01 – 0.01] ppm interval. If the mean of the bias was within this interval, the difference between the corresponding pair of QSM pipelines was considered negligible.

A bias greater than |0.01| ppm was observed for the NLFit pipeline relative to all other pipelines in the CN, and StrS; for the Susc-TGV-wAvg pipeline relative to all other pipelines in the GP, and relative to the TE-wAvg and Susc-wAvg pipelines in the StrS (Figure 7). These results suggests that, for accurate *χ* quantification, some of the significant differences detected by the sign test, e.g., between the TE-wAvg, SNR-wAvg and Susc-wAvg pipelines, may be negligible (Figures 3B and 7).

## Discussion

Aiming to elucidate the optimal strategy for multi-echo combination for QSM, this study compared multi-echo combination methods applied at different stages of the QSM processing pipeline before or after LBMs for phase unwrapping or background field removal. Each pipeline was applied to numerically simulated data and images from healthy volunteers.

The higher relative accuracy of the NLFit pipeline in the numerical phantom simulations and *in vivo* suggests that, for QSM, combining the temporally unwrapped multi-echo phase before applying LBMs for spatial phase unwrapping or Δ*B_Bg_* removal is preferable to averaging the TE-dependent LBM-processed phase or *χ*. This suggestion appears to conflict with the higher RMSEs associated with the NLFit pipeline compared to other pipelines in the phantom simulations (Figure 2 and Supplementary Information Figure S3). However, the RMSE jointly reflects systematic and random errors, as it measures the bias between the estimated and true value, but also reflects the variability of the estimated *χ* relative to its average value (3). Thus, the RMSE must always be interpreted in combination with complementary measurements of bias and precision. Furthermore, RMSEs of Δ*B_Loc_* are difficult to interpret. In contrast with RMSEs of *χ*, they allow to compare different pipelines without the effect of Δ*B_Loc_*-to-*χ* inversion. However, they are voxel-based measures based on a signal that is intrinsically nonlocal, because Δ*B_Loc_* variations extend beyond the anatomical region of *χ* shift that generated them (19). Thus, the best set of metrics for comparing images generated by a processing pipeline for QSM is still an active area of research (3,4).

There are several potential explanations as to why LBMs applied before multi-echo combination reduce the overall accuracy of QSM. Firstly, in contrast with path-based or region-growing-based phase unwrapping, Laplacian phase unwrapping usually removes some Δ*B_Bg_* components from the input phase image (30). Thus, the consistently lower accuracy of *χ* calculated using the TE-wAvg processing pipeline was probably driven by the incorrect assumption that the TE-dependent LBM-unwrapped phase corresponded to the true unwrapped phase. LBMs for Δ*B_Bg_* removal also applied tSVD (with a larger truncation threshold) but here the high pass filtering effect was expected, as background fields are slowly varying. Finally, the similar accuracy and values of *χ* calculated using the TE-wAvg, SNR-wAvg and Susc-wAvg pipelines suggest a negligible difference between averaging the Laplacian unwrapped phase over TEs before (TE-wAvg pipeline) or after TE-dependent Laplacian Δ*B_Bg_* removal (SNR-wAvg and Susc-wAvg pipelines).

In the numerical phantom simulations, all processing pipelines resulted in higher SDs of *χ* compared to the ground-truth, suggesting that noise in the multi-echo signal phase was amplified by all pipelines. This result is in line with the known noise amplification of ill-posed inverse problems. However, the estimated SDs of *χ* varied across pipelines. In the simulations, the Susc-TGV-wAvg pipeline had the smallest SDs of *χ* (Figure 3A). However, *in vivo*, the NLFit pipeline generally had the smallest SDs of *χ* (Figure 3B). Both the TGV reconstruction pipeline and shortcomings of the numerically simulated data could explain these discrepancies in the performance of the Susc-TGV-wAvg pipeline. The numerically simulated data were generated based on a digital phantom which, despite varying regional *χ* values in a realistic fashion (see Supporting Information) ultimately still appeared as a smooth piece-wise constant model (Figures 2A, G). As previously observed (3), piece-wise constant geometries allow good recovery of the underlying *χ* distribution using TGV-based algorithms, because the piece-wise constant constraints exactly match the underlying *χ* distribution. However, in regions with flow, anisotropic *χ* distributions, or microstructure, these numerical models are likely to depart from a realistic representation of the tissue *χ*. To overcome the limitations of this assumption, future studies could exploit a newly developed realistic head phantom for QSM, which does not have a piecewise constant *χ* distribution and incorporates microstructural effects (50).

In both numerical simulations and healthy volunteers, based on the line profiles traced on the noise maps and streaking artifact reduction, the NLFit pipeline had better noise mitigation (Figures 5 and 6). This result suggests that combining the temporally unwrapped multi-echo phase by nonlinear complex fitting, designed to account for noise in the complex signal (12), results in better noise management than combining the multi-echo phase by averaging. As previously shown (12), errors in the combined field map mainly result from both noise in the signal phase and phase unwrapping errors near high-*χ* regions (e.g., the veins). Both sources of error were successfully managed by nonlinear complex fitting, as shown by the dramatic reduction of streaking artifacts in Figure 6A. In line with previous observations (51), this result also suggests that the regularization strategy employed by the Δ*B_Loc_*-to *χ* step can mitigate artifactual streaking errors only when major sources of error in the field map have been tightly constrained. At visual inspection *in vivo*, the SNR-wAvg was the second-best pipeline for the mitigation of streaking artifacts (Figure 6). Thus, if applying nonlinear complex fitting is not possible, averaging using the SNR-weighting-based method could offer the best alternative for noise reduction.

All these indications do not necessarily apply to *χ* estimation in WM tissue. Indeed, due to WM’s ordered microstructure, a comprehensive estimation of *χ* in WM requires acquiring GRE images at multiple head orientations and modelling *χ* as a tensor (52,53). Thus, further studies are needed to evaluate the applicability of these results to WM tissue.

In the present study, all experiments were limited to one field strength (i.e., 3 T). As tissue relaxation times (e.g., 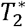) shorten with increasing field strength but the signal phase at a given TE increases, further work is needed to assess the relevance of these results at ultrahigh fields. Finally, it must be noted that in the numerical phantom simulations, all processing pipelines underestimated *χ^True^* (Figure 3). However, in QSM, some degree of underestimation is always expected, because of the ill-posed nature of the Δ*B_Loc_*-to-*χ* inverse problem (20).

## Conclusions

The higher accuracy of regional *χ* values and better noise management of the NLFit pipeline suggest that, for QSM, combining the multi-echo phase by nonlinearly fitting over TEs before applying LBMs is preferable to combining the TE-dependent LBM-processed phase or *χ* by averaging.

## Data availability statement

The Matlab code used to run the analyses in this study is available at https://github.com/emmabiondetti/multi-echo-qsm. The repository’s main page lists all dependencies on code developed by other groups.

## Acknowledgments

We thank all the healthy volunteers who participated in this study. We also thank Dr. Rosa Cortese and Dr. Floriana De Angelis (Queen Square MS Centre, UCL Queen Square Institute of Neurology, UCL, London, UK) for their help with the MRI scans. We are indebted to Prof. Claudia Gandini Wheeler-Kingshott (NMR Unit, Queen Square MS Centre, UCL Queen Square Institute of Neurology, UCL, London, UK) for her support with the MRI scans performed at the Queen Square MS Centre, UCL.

**Supporting Information Figure S1.** Properties of the numerical phantom. *M*_0_ in arbitrary units (a. u.), 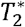 in ms and *χ* in parts per million (ppm) assigned to various ROIs in the numerical phantom are shown in **(A)**. The location of these ROIs is shown in the *χ* **(B)** and T_2_ maps **(C)** of the numerical phantom.

**Supporting Information Table S2.** Total image processing time. The table shows the time required to run each pipeline for the numerical phantom simulations and data acquired *in vivo*. The time reported for the SNR-wAvg pipeline does not include the time required for 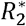 mapping, as optimizing this step was outside the scope of the present study. For the Susc-TGV-wAvg pipeline, approximate timings are reported because image reconstruction was performed using first Neurodesk (QSM calculation at each TE) and then Matlab (multi-echo combination).

**Supporting Information Figure S3.** Δ*B_Loc_* maps calculated using distinct multi-echo combination methods in the numerical phantom simulations. The same transverse and sagittal slices are shown for the ground-truth local field map **(A, G)**, and for the local field maps calculated using NLFit **(B, H)**, TE-wAvg **(C, I)**, SNR-wAvg **(D, J)**, and Susc-wAvg at each TE **(E, O)**. The figure also shows the difference between each local field map and the ground truth **(P-E2)**. The bottom row shows the root mean squared errors (RMSEs) of Δ*B_Loc_* for each pipeline.

## Notes

**Funding information**, E. Biondetti was supported by the UK Engineering and Physical Sciences Research Council (EPSRC) (award number: 1489882). A. Karsa was supported by the EPSRC-funded UCL Centre for Doctoral Training in Medical Imaging (EP/L016478/1) and the Department of Health’s National Institute for Health Research funded Biomedical Research Centre at University College London Hospitals. D. L. Thomas was supported by the UCL Leonard Wolfson Experimental Neurology Centre (PR/ylr/18575). The Queen Square MS Centre, where part of the MRI scans for this work were performed, is supported by grants from the UK MS Society and by the National Institute for Health Research University College London Hospitals Biomedical Research Centre. F. Grussu was supported by PREdICT, a study at the Vall d’Hebron Institute of Oncology in Barcelona funded by AstraZeneca, and currently receives funding from the postdoctoral fellowships program Beatriu de Pinós (2020 BP 00117), funded by the Secretary of Universities and Research (Government of Catalonia).

### Competing Interest Statement

The authors have declared no competing interest.

### Summary of Updates

The manuscript title has been updated. Significant revisions to the text structure have been made for improved clarity.

