## Supporting Information for "Multi-echo Quantitative Susceptibility Mapping: How to Combine Echoes for Accuracy and Precision at 3 T"

### Biondetti et al. Supporting Information

#### Noise in total field map calculated by fitting

Regressing the phase on TE using a weighted linear least-square fit enables the calculation of a field map ($\omega^{Fit}=\gamma\Delta B_{Tot}^{Fit}$ in rad/s) and a phase offset ($\phi_{0}^{Fit}$) by minimizing the following weighted residual sum of squares (WRSS):

$$\begin{aligned} WRSS\left( \boldsymbol{r} \right)=\sum_{i=1}^{n} w_{i}\left( \phi\left( \boldsymbol{r}, TE_{i} \right)-\left( \omega^{Fit}\left( \boldsymbol{r} \right)TE_{i}+\phi_{0}^{Fit}\left( \boldsymbol{r} \right) \right) \right)^{2}\#\left( S1 \right) \end{aligned}$$

Where $w_{i}$ is a weighting factor equal to the inverse variance of the phase, i.e.$w_{i}=1/\sigma^{2}\left( \phi\left( TE_{i} \right) \right)$ and $n$ is the total number of TEs (1).

The values of $\omega^{Fit}$ and $\phi_{0}^{Fit}$ that jointly minimize Equation S1 are calculated by jointly setting to zero the partial derivatives of WRSS relative to $\omega^{Fit}$ and $\phi_{0}^{Fit}$:

$$\begin{aligned} \left\{ \begin{aligned} \frac{\partial WRSS(\boldsymbol{r})}{\partial\omega^{Fit}}=0 \\ \frac{\partial WRSS(\boldsymbol{r})}{\partial\phi_{0}^{Fit}}=0 \end{aligned} \right.\#\left( S2 \right) \end{aligned}$$

Omitting the $\boldsymbol{r}$ dependency (because fitting is performed voxel by voxel), solving the system in Equation S2 yields (2):

$$\begin{aligned} \omega^{Fit}=\frac{\sum_{i=1}^{n} w_{i}\sum_{i=1}^{n} w_{i}TE_{i}\phi\left( TE_{i} \right)-\sum_{i=1}^{n} w_{i}\phi\left( TE_{i} \right)\sum_{i=1}^{n} w_{i}TE_{i}}{\sum_{i=1}^{n} w_{i}\sum_{i=1}^{n} w_{i}TE_{i}^{2}-\left( \sum_{i=1}^{n} w_{i}TE_{i} \right)^{2}}\#\left( S3 \right) \end{aligned}$$

Equation 14 in the main manuscript is derived from Equation S3 and error propagation.

#### Noise propagation from the local field to the susceptibility map

Based on Equation 3 in the main manuscript and the circular convolution theorem, the susceptibility map $\chi$ can be calculated as:

$$\begin{aligned} \chi\left( \boldsymbol{r} \right)=\frac{1}{B_{0}}FT^{-1}\left\{ D_{Tik}^{-1}\left( \boldsymbol{k} \right)FT\left( \Delta B_{Loc}\left( \boldsymbol{r} \right) \right) \right\}\#\left( S4 \right) \end{aligned}$$

where $D_{Tik}$ is the Tikhonov-regularized dipole kernel in k-space, $\Delta B_{Loc}$ is the local field in image space, $B_{0}$ is the static magnetic field strength, $FT$ and $FT^{-1}$ respectively denote the direct and inverse Fourier transforms, and $\boldsymbol{r}$ and $\boldsymbol{k}$ respectively denote coordinates in image and Fourier space. The Fourier transform of the local field map is:

$$\begin{aligned} FT\left\{ \Delta B_{Loc}\left( \boldsymbol{r} \right) \right\}=\int_{\boldsymbol{r}} \Delta B_{Loc}\left( \boldsymbol{r} \right)\exp\left( -i2\pi\boldsymbol{kr} \right)d\boldsymbol{r}\#\left( S5 \right) \end{aligned}$$

Based on Equations S4 and S5, the susceptibility can be written as:

$$\begin{aligned} \chi\left( \boldsymbol{r} \right)=\frac{1}{B_{0}}\int_{\boldsymbol{k}} D^{-1}\left( \boldsymbol{k} \right)\left( \int_{\boldsymbol{s}} \Delta B_{Loc}\left( \boldsymbol{s} \right)\exp\left( -i2\pi\boldsymbol{ks} \right)d\boldsymbol{s} \right)\exp\left( i2\pi\boldsymbol{kr} \right)d\boldsymbol{k}\#\left( S6 \right) \end{aligned}$$

where $\boldsymbol{s}$ is an auxiliary variable of integration in image space. It must be noted that here the Fourier transform is expressed as an integral operation. However, because this operation is performed on the discrete image domain, the integral actually denotes a discrete Fourier transform.

The error propagation from the local field to the susceptibility map is calculated as:

$$\sigma^{2}\left( \chi\left( \boldsymbol{r} \right) \right)=\int_{\boldsymbol{r}} \left( \frac{\partial\chi\left( \boldsymbol{r} \right)}{\partial\Delta B_{Loc}\left( \boldsymbol{r} \right)} \right)^{2}\sigma^{2}\left( \Delta B_{Loc}\left( \boldsymbol{r} \right) \right)d\boldsymbol{r}=$$

$$\begin{aligned} \int_{\boldsymbol{r}} \left( \frac{\partial}{\partial\Delta B_{Loc}\left( \boldsymbol{r} \right)}\left( \frac{1}{B_{0}}\int_{\boldsymbol{k}} D^{-1}\left( \boldsymbol{k} \right)\left( \int_{\boldsymbol{s}} \Delta B_{Loc}\left( \boldsymbol{s} \right)\exp\left( -i2\pi\boldsymbol{ks} \right)d\boldsymbol{s} \right)\exp\left( i2\pi\boldsymbol{k}\boldsymbol{r}^{\boldsymbol{'}} \right)d\boldsymbol{k} \right) \right)^{2}\sigma^{2}\left( \Delta B_{Loc}\left( \boldsymbol{r} \right) \right)d\boldsymbol{r}\#\left( S7 \right) \end{aligned}$$

where $\boldsymbol{r}^{\boldsymbol{'}}$ is an auxiliary variable of integration in image space. Owing to the Leibniz integral rule, the partial derivative can be moved under the integral operator giving:

$$\begin{aligned} \sigma^{2}\left( \chi\left( \boldsymbol{r} \right) \right)= \\ \int_{\boldsymbol{r}} \left( \frac{1}{B_{0}}\int_{\boldsymbol{k}} D^{-1}\left( \boldsymbol{k} \right)\left( \int_{\boldsymbol{s}} \frac{\partial\Delta B_{Loc}\left( \boldsymbol{s} \right)\exp\left( -i2\pi\boldsymbol{ks} \right)}{\partial\Delta B_{Loc}\left( \boldsymbol{r} \right)}d\boldsymbol{s} \right)\exp\left( i2\pi\boldsymbol{k}\boldsymbol{r}^{\boldsymbol{'}} \right)d\boldsymbol{k} \right)^{2}\sigma^{2}\left( \Delta B_{Loc}\left( \boldsymbol{r} \right) \right)d\boldsymbol{r}\#\left( S8 \right) \end{aligned}$$

The partial derivative operation in Equation S8 can be written as:

$$\begin{aligned} \frac{\partial\Delta B_{Loc}\left( \boldsymbol{s} \right)}{\partial\Delta B_{Loc}\left( \boldsymbol{r} \right)}=\delta\left( \boldsymbol{s}-\boldsymbol{r} \right)=\left\{ \begin{aligned} 1, \boldsymbol{s}=\boldsymbol{r} \\ 0, \boldsymbol{s}\neq\boldsymbol{r} \end{aligned} \right.\#\left( S9 \right) \end{aligned}$$

Where $\delta\left( \boldsymbol{s}-\boldsymbol{r} \right)$ denotes a Dirac delta function, whose center has been spatially shifted from $\boldsymbol{s}$ to $\boldsymbol{r}$. The physical motivation for this shift is that $\Delta B_{Loc}$ results from a spatial convolution operation between the underlying tissue magnetic susceptibility $\chi$ and the unit magnetic dipole. Thus, Equation S8 can be written as:

$$\begin{aligned} \sigma^{2}\left( \chi\left( \boldsymbol{r} \right) \right)=\int_{\boldsymbol{r}} \left( \frac{1}{B_{0}}\int_{\boldsymbol{k}} D^{-1}\left( \boldsymbol{k} \right)\left( \int_{\boldsymbol{s}} \delta\left( \boldsymbol{s}-\boldsymbol{r} \right)\exp\left( -i2\pi\boldsymbol{ks} \right)d\boldsymbol{s} \right)\exp\left( i2\pi\boldsymbol{k}\boldsymbol{r}^{\boldsymbol{'}} \right)d\boldsymbol{k} \right)^{2}\sigma^{2}\left( \Delta B_{Loc}\left( \boldsymbol{r} \right) \right)d\boldsymbol{r}\#= \end{aligned}$$

$$=\int_{\boldsymbol{r}} \left( \frac{1}{B_{0}}\int_{\boldsymbol{k}} D^{-1}\left( \boldsymbol{k} \right)\exp\left( -i2\pi\boldsymbol{kr} \right)\exp\left( i2\pi\boldsymbol{k}\boldsymbol{r}^{\boldsymbol{'}} \right)d\boldsymbol{k} \right)^{2}\sigma^{2}\left( \Delta B_{Loc}\left( \boldsymbol{r} \right) \right)d\boldsymbol{r=}$$

$$\begin{aligned} =\int_{\boldsymbol{r}} \left( \frac{1}{B_{0}}\int_{\boldsymbol{k}} D^{-1}\left( \boldsymbol{k} \right)\exp\left( i2\pi\boldsymbol{k}\left( \boldsymbol{r}^{\boldsymbol{'}}\boldsymbol{-r} \right) \right)d\boldsymbol{k} \right)^{2}\sigma^{2}\left( \Delta B_{Loc}\left( \boldsymbol{r} \right) \right)d\boldsymbol{r}\#\left( S10 \right) \end{aligned}$$

In Equation S10, the integral in $d\boldsymbol{k}$ is the Fourier transform of the inverse unit magnetic dipole $d_{z}^{-1}$ evaluated in $\boldsymbol{r}^{\boldsymbol{'}}\boldsymbol{-r}$:

$$\sigma^{2}\left( \chi\left( \boldsymbol{r} \right) \right)=\int_{\boldsymbol{r}} \left( \frac{1}{B_{0}}d_{z}^{-1}\left( \boldsymbol{r}^{\boldsymbol{'}}-\boldsymbol{r} \right) \right)^{2}\sigma^{2}\left( \Delta B_{Loc}\left( \boldsymbol{r} \right) \right)d\boldsymbol{r}=\frac{1}{B_{0}^{2}}\int_{\boldsymbol{r}} \left( d_{z}^{-1}\left( \boldsymbol{r}^{\boldsymbol{'}}-\boldsymbol{r} \right) \right)^{2}\sigma^{2}\left( \Delta B_{Loc}\left( \boldsymbol{r} \right) \right)d\boldsymbol{r}$$

$$\begin{aligned} =\frac{1}{B_{0}^{2}}d_{z}^{-2}(\boldsymbol{r})\star\sigma^{2}\left( \Delta B_{Loc}\left( \boldsymbol{r} \right) \right)\#\left( S11 \right) \end{aligned}$$

where $\star$ denotes a convolution operation. Equation 20 in the main manuscript is obtained as the square root of Equation S11. To conclude, the noise affecting the susceptibility map calculated from a combined field map is a function of the convolution between the variance of the corresponding local field map and the (regularized) inverse dipole squared.

#### Noise propagation in the Susc-wAvg pipeline

Noise propagation into the combined susceptibility map calculated using the Susc-wAvg pipeline can be calculated as follows:

$$\begin{aligned} \sigma^{\left( \chi^{Susc-wAvg} \right)}=\left( \frac{1}{\left( B_{0}\gamma n \right)^{2}}\sum_{i=1}^{n} \sigma^{2}\left( \frac{A_{i}}{B}C_{i} \right) \right)^{\frac{1}{2}}\#\left( S12 \right) \end{aligned}$$

where	$A_{i}=M^{2}\left( TE_{i} \right)TE_{i}^{2}$	$B=\sum_{j=1}^{n} M^{2}\left( TE_{j} \right)TE_{j}^{2}$	$C_{i}=\chi\left( TE_{i} \right)$

$$\begin{aligned} \sigma^{2}\left( \frac{A_{i}}{B} \right)=\frac{\sigma^{2}\left( A_{i} \right)}{B^{2}}+\frac{A_{i}^{2}}{B^{4}}\sigma^{2}\left( B \right)-\frac{2A_{i}}{B^{3}}cov(A_{i},B)\#\left( S13 \right) \end{aligned}$$

where $cov\left( \cdot\right)$ denotes the covariance. Based on the linearity of covariances, this term is calculated as:

$$\begin{aligned} cov\left( A_{i},B \right)=cov\left( M^{2}\left( TE_{i} \right)TE_{i}^{2},\sum_{j=1}^{n} M^{2}\left( TE_{j} \right)TE_{j}^{2} \right)= \\ =\sum_{j=1}^{n} cov\left( M^{2}\left( TE_{i} \right)TE_{i}^{2},M^{2}\left( TE_{j} \right)TE_{j}^{2} \right)\#\left( S14 \right) \end{aligned}$$

By assuming temporally uncorrelated magnitude signals, the covariance term in Equation S14 can be written as:

$$\begin{aligned} cov\left( A_{i},B \right)=\left\{ \begin{aligned} \sigma^{2}(A_{i}), i=j \\ 0, i\neq j \end{aligned} \right.\#\left( S15 \right) \end{aligned}$$

Then:

$$\begin{aligned} \sigma^{2}\left( \frac{A_{i}}{B} \right)=\frac{\sigma^{2}\left( A_{i} \right)}{B^{2}}+\frac{A_{i}^{2}}{B^{4}}\sigma^{2}\left( B \right)-\frac{2A_{i}}{B^{3}}\sigma^{2}(A_{i})\#\left( S16 \right) \end{aligned}$$

$$\begin{aligned} \sigma^{2}\left( \frac{A_{i}}{B}C_{i} \right)=C_{i}^{2}\sigma^{2}\left( \frac{A_{i}}{B} \right)+\left( \frac{A_{i}}{B} \right)^{2}\sigma^{2}\left( C_{i} \right)+\frac{{2C}_{i}A_{i}}{B}cov\left( C_{i},\frac{A_{i}}{B} \right)\#\left( S17 \right) \end{aligned}$$

where $cov\left( C_{i},A_{i}/B \right)=0$ because we can assume that the two variables are uncorrelated. Thus:

$$\begin{aligned} \sigma^{2}\left( \frac{A_{i}}{B}C_{i} \right)=C_{i}^{2}\left( \frac{\sigma^{2}\left( A_{i} \right)}{B^{2}}+\frac{A_{i}^{2}}{B^{4}}\sigma^{2}\left( B \right)-\frac{2A_{i}}{B^{3}}\sigma^{2}\left( A_{i} \right) \right)+\left( \frac{A_{i}}{B} \right)^{2}\sigma^{2}\left( C_{i} \right)\#\left( S18 \right) \end{aligned}$$

$$\begin{aligned} \sigma^{2}\left( A_{i} \right)=\sigma^{2}\left( M^{2}\left( TE_{i} \right)TE_{i}^{2} \right)=TE_{i}^{4}\sigma^{2}\left( M^{2}\left( TE_{i} \right) \right)=4M^{2}\left( TE_{i} \right)TE_{i}^{4}\sigma^{2}\left( M\left( TE_{i} \right) \right)\#\left( S19 \right) \end{aligned}$$

$$\begin{aligned} \sigma^{2}\left( B \right)= \sum_{j=1}^{n} \sigma^{2}\left( M^{2}\left( TE_{j} \right)TE_{j}^{2} \right)=\sum_{j=1}^{n} 4M^{2}\left( TE_{j} \right)TE_{j}^{4}\sigma^{2}\left( M\left( TE_{j} \right) \right)\#\left( S20 \right) \end{aligned}$$

$$\sigma^{2}\left( \chi^{Susc-wAvg} \right)==\frac{1}{\left( B_{0}\gamma n \right)^{2}}\sum_{i=1}^{n} \chi^{2}\left( TE_{i} \right)\left( \frac{4M^{2}\left( TE_{i} \right)TE_{i}^{4}\sigma^{2}\left( M\left( TE_{i} \right) \right)}{\sum_{j=1}^{n} \left( M^{2}\left( TE_{j} \right)TE_{j}^{2} \right)^{2}}+\frac{M^{4}\left( TE_{i} \right)TE_{i}^{4}}{\sum_{j=1}^{n} \left( M^{2}\left( TE_{j} \right)TE_{j}^{2} \right)^{4}}\sum_{j=1}^{n} 4M^{2}\left( TE_{j} \right)TE_{j}^{4}\sigma^{2}\left( M\left( TE_{j} \right) \right)-\frac{2M^{2}\left( TE_{i} \right)TE_{i}^{2}}{\left( \sum_{j=1}^{n} M^{2}\left( TE_{j} \right)TE_{j}^{2} \right)^{3}}4M^{2}\left( TE_{i} \right)TE_{i}^{4}\sigma^{2}\left( M\left( TE_{i} \right) \right) \right)+\left( \frac{M^{2}\left( TE_{i} \right)TE_{i}^{2}}{\sum_{j=1}^{n} M^{2}\left( TE_{j} \right)TE_{j}^{2}} \right)^{2}\sigma^{2}\left( \chi\left( TE_{i} \right) \right)$$

$$(S21)$$

The noise in the Susc-wAvg susceptibility map is obtained as the square root of Equation S21.

#### Numerical phantom simulations

Realistic SD values were calculated by multiplying the corresponding ground-truth mean of $\chi$, $M_{0}$ or $T_{2}^{*}$ by realistic coefficients of variation (CV) of the mean. These were calculated as the ratio of the SD to the mean based on measurements in previous studies. Particularly, a CV = 4% for $T_{2}^{*}$ measurements was determined based on Yao et al. (3); a CV = 1% for $M_{0}$ was determined based on Cordes et al. (4); a CV = 20% for $\chi$ was determined based on the average CV calculated by Liu et al. using the COSMOS method (calculation of susceptibility using multiple orientation sampling) (5).

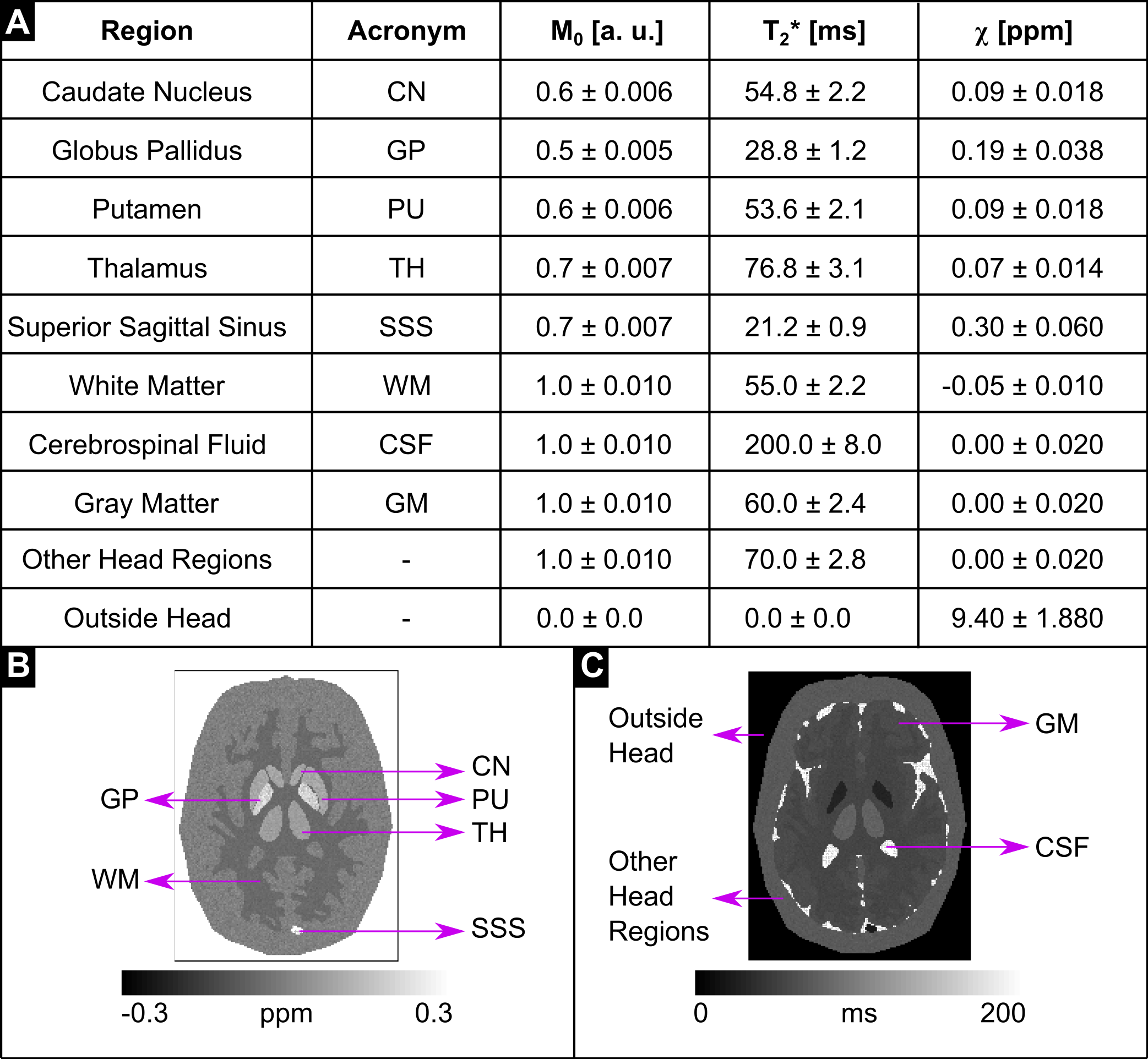

**Supporting Information Figure S1.** Properties of the numerical phantom. $M_{0}$ in arbitrary units (a. u.), $T_{2}^{*}$ in ms and $\chi$ in parts per million (ppm) assigned to various ROIs in the numerical phantom are shown in **(A)**. The location of these ROIs is shown in the $\chi$ **(B)** and $T_{2}^{*}$ maps **(C)** of the numerical phantom.

|  | **NLFit** | **TE-wAvg** | **SNR-wAvg** | **Susc-wAvg** | **Susc-TGV-wAvg** |
| --- | --- | --- | --- | --- | --- |
| Numerical phantom simulations | 5 min 4 s | 3 min 46 s | 3 min 56 s | 16 min 39 s | Approx. 75 min |
| Healthy volunteers | 2 min 22 s | 1 min 11 s | 1 min 20 s | 5 min 16 s | Approx. 35 min |

**Supporting Information Table S2.** Total image processing time. The table shows the time required to run each pipeline for the numerical phantom simulations and data acquired *in vivo*. The time reported for the SNR-wAvg pipeline does not include the time required for $R_{2}^{*}$ mapping, as optimizing this step was outside the scope of the present study. For the Susc-TGV-wAvg pipeline, approximate timings are reported because image reconstruction was performed using first Neurodesk (QSM calculation at each TE) and then Matlab (multi-echo combination).

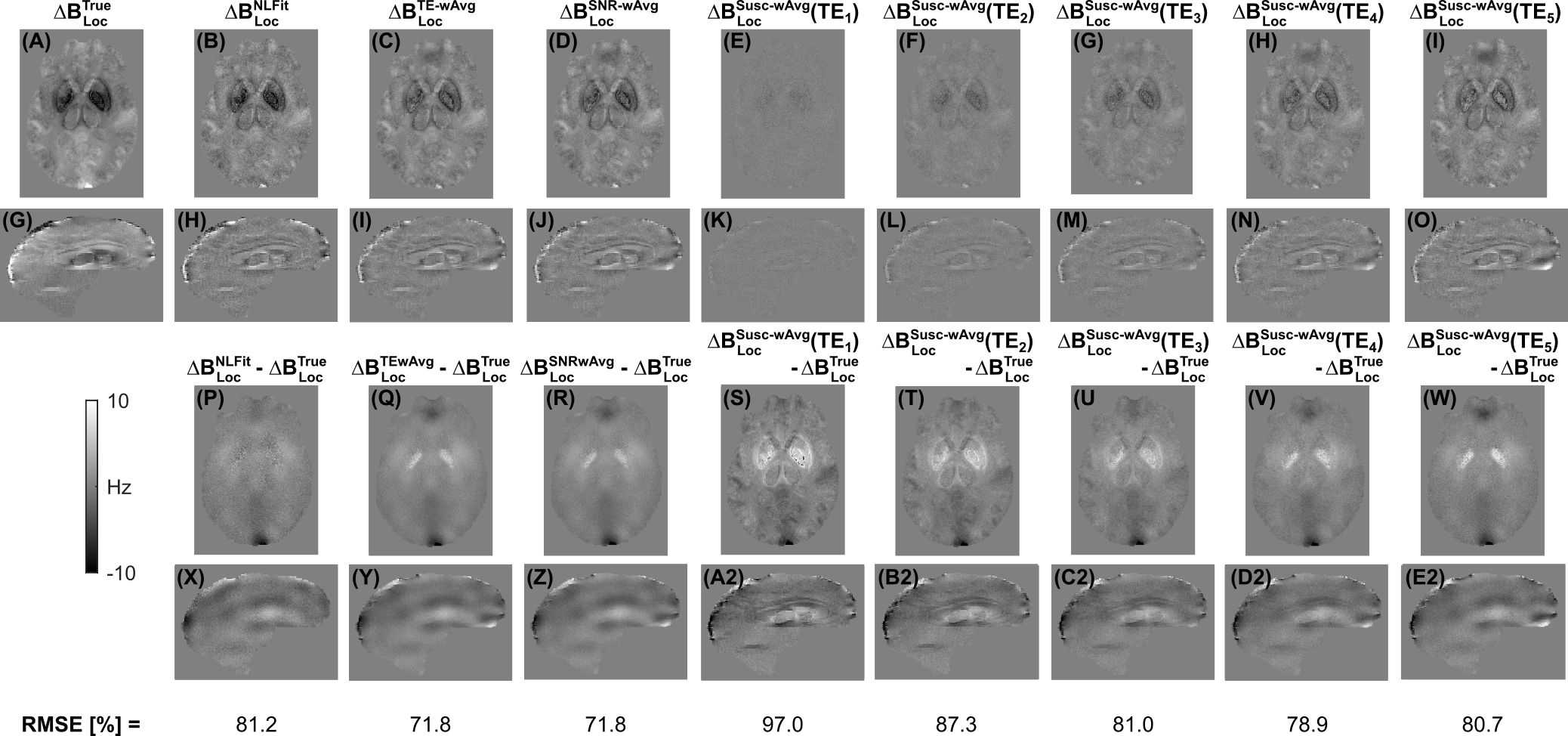

**Supporting Information Figure S3.** $\Delta B_{Loc}$ maps calculated using distinct multi-echo combination methods in the numerical phantom simulations. The same transverse and sagittal slices are shown for the ground-truth local field map **(A, G)**, and for the local field maps calculated using NLFit **(B, H)**, TE-wAvg **(C, I)**, SNR-wAvg **(D, J)**, and Susc-wAvg at each TE **(E, O)**. The figure also shows the difference between each local field map and the ground truth **(P-E2)**. The bottom row shows the root mean squared errors (RMSEs) of $\Delta B_{Loc}$ for each pipeline.
